# Association Between Dietary Vitamin B1 and the Risk of Endometriosis: A Cross-Sectional Study Based on NHANES 2003–2006

**DOI:** 10.1101/2025.02.21.639433

**Authors:** Jing Zhang, Huiwen Guo, Zhigang Zhou

## Abstract

Endometriosis is a chronic gynecological condition influenced by inflammation and oxidative stress. Dietary vitamin B1 plays a role in reducing oxidative stress, which may influence endometriosis risk. However, limited research explores this association.

**Objective:** To investigate the association between dietary vitamin B1 intake and the risk of endometriosis, using NHANES 2003–2006 data. Vitamins B2 and B6 are included as reference comparisons.

**Methods:** Data from 2,462 participants in NHANES 2003–2006 were analyzed. Logistic regression models were used to examine the association between vitamin B1 intake and endometriosis risk, with adjustments for covariates. Subgroup analyses and restricted cubic spline (RCS) regressions were applied to assess non-linear relationships. Vitamins B2 and B6 served as reference nutrients for comparison.

**Results:** Vitamin B1: Higher intake of dietary vitamin B1 was significantly associated with reduced risk of endometriosis (OR = 0.76, 95% CI: 0.58– 0.99, P = 0.044 in Model 4).

Vitamin B2 and B6: No significant associations were observed (P > 0.05).

RCS regression demonstrated a non-linear relationship for vitamin B1, with a protective effect observed at intake levels > 1.84 mg/day.

**Conclusion:** Higher dietary vitamin B1 intake is associated with a reduced risk of endometriosis. While B2 and B6 showed no significant associations, their trends provide additional insight for future research.

## Introduction

Endometriosis is a common, chronic inflammatory gynecological condition characterized by the ectopic growth of endometrial-like tissue outside the uterine cavity. It affects approximately 10% of women of reproductive age globally, leading to symptoms such as pelvic pain, dysmenorrhea, and infertility, significantly impairing the quality of life ^1^; ^2^. The pathogenesis of endometriosis remains complex and multifactorial, involving genetic predisposition, hormonal imbalances, immune dysregulation, and oxidative stress ^3^; ^4^.

The Role of Oxidative Stress and Inflammation in Endometriosis Oxidative stress, defined as an imbalance between reactive oxygen species (ROS) production and antioxidant defenses, plays a pivotal role in the development and progression of endometriosis. Elevated ROS levels contribute to peritoneal inflammation, angiogenesis, and tissue adhesion, exacerbating the ectopic implantation and growth of endometrial tissue ^5^; ^6^. This oxidative environment not only enhances cellular damage but also perpetuates chronic inflammation, a hallmark of endometriosis ^7^.

### Potential Role of B Vitamins in Endometriosis

Dietary factors have emerged as modifiable risk factors for endometriosis, with increasing evidence highlighting the role of micronutrients in mitigating oxidative stress and inflammation. Among these, B vitamins, particularly thiamine (vitamin B1), riboflavin (vitamin B2), and pyridoxine (vitamin B6), are essential coenzymes involved in energy metabolism, cellular repair, and redox homeostasis.

Vitamin B1 (Thiamine): Thiamine plays a crucial role in the pentose phosphate pathway, which generates NADPH, a key molecule in antioxidant defense. Adequate thiamine intake enhances mitochondrial function, reduces ROS production, and suppresses inflammation ^8^. Given that oxidative stress is elevated in the peritoneal cavity of women with endometriosis, vitamin B1’s role in reducing oxidative damage may be particularly relevant ^9^.

Vitamin B2 (Riboflavin): Riboflavin acts as a cofactor for glutathione reductase, a key antioxidant enzyme, and has been shown to reduce systemic oxidative stress ^10^. However, studies exploring its association with endometriosis are limited.

Vitamin B6 (Pyridoxine): Vitamin B6 exhibits anti-inflammatory properties by modulating cytokine production and supporting homocysteine metabolism, a process implicated in oxidative stress and inflammation ^11^.

Despite their biological plausibility, evidence linking dietary B-vitamin intake to endometriosis remains sparse. A few observational studies suggest that higher intake of certain B vitamins may reduce the risk of inflammatory and oxidative stress-related diseases, including cardiovascular disease and polycystic ovary syndrome (PCOS) ^12^; ^13^. However, specific data on vitamin B1 and its association with endometriosis are lacking.

### Rationale for the Study

Given the critical role of oxidative stress and inflammation in the pathophysiology of endometriosis, understanding the potential protective effects of vitamin B1 is of significant importance. Additionally, examining vitamins B2 and B6 as reference nutrients provides a comparative context to elucidate the unique contribution of thiamine. To address this gap, we conducted a cross-sectional analysis using data from the National Health and Nutrition Examination Survey (NHANES) 2003– 2006. This study aims to investigate the association between dietary vitamin B1 intake and the risk of endometriosis, while including vitamins B2 and B6 as comparative references.

## Methods

### Study Design and Data Source

This study utilized a cross-sectional design based on publicly available data from the National Health and Nutrition Examination Survey (NHANES) 2003–2006. NHANES, conducted by the National Center for Health Statistics (NCHS), employs a multistage stratified probability sampling design to generate nationally representative health and nutritional data of the U.S. population. The study protocol was approved by the NCHS Research Ethics Review Board, and all participants provided informed consent.

### Study Population

From a total of 20,470 participants in the NHANES 2003–2006 dataset, we applied the following inclusion and exclusion criteria (Figure 1):

**Figure 1.**
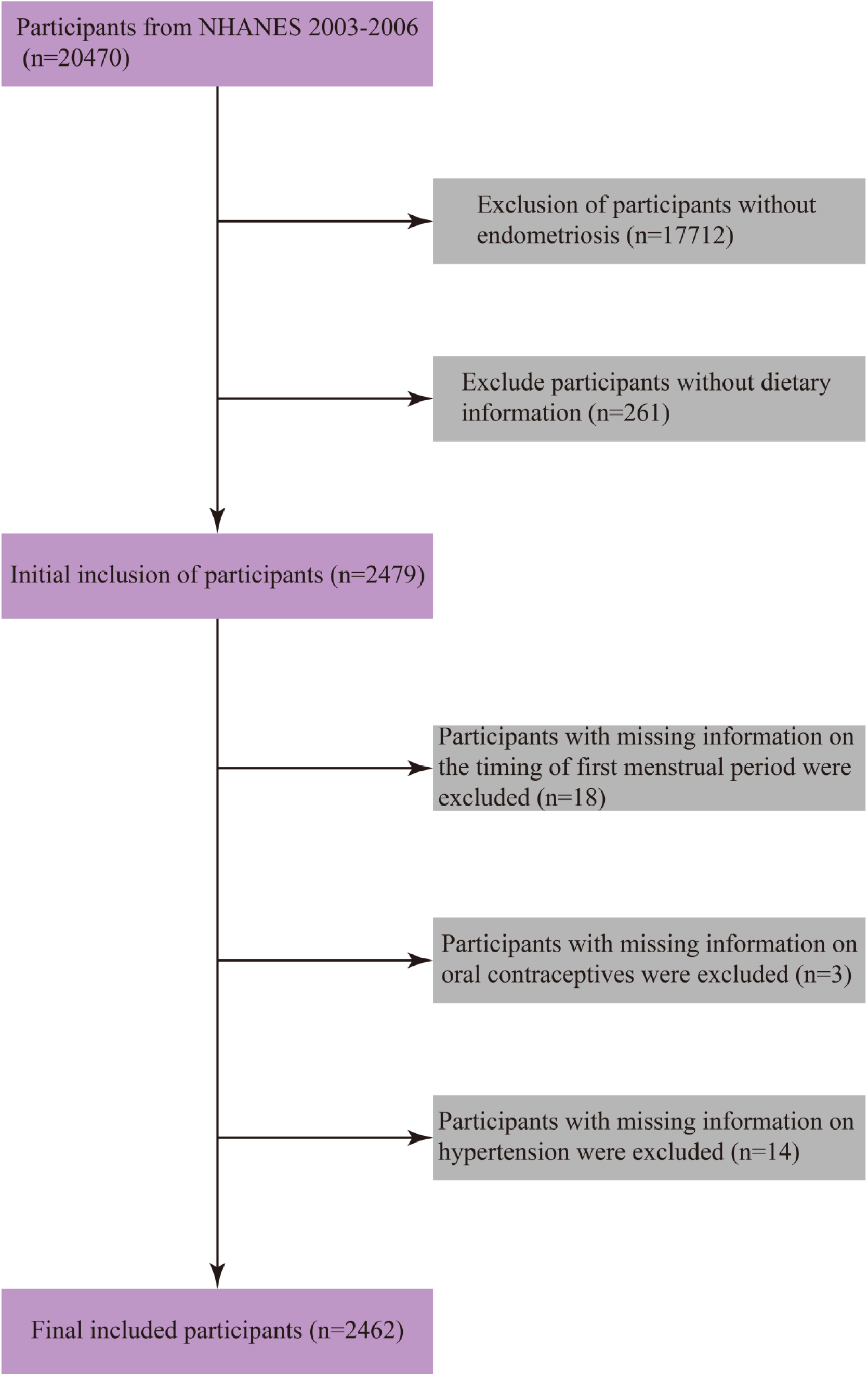
Flowchart of the study population

Exclusion of participants without endometriosis diagnosis data (n = 17,712).

Exclusion of participants missing dietary information on vitamin B intake (n = 261).

Exclusion of participants with missing key covariates: menarche age (n = 18), oral contraceptive use (n = 3), and hypertension status (n = 14).

After exclusions, the final analytical sample included 2,462 participants (Figure 1).

### Exposure Assessment: Dietary Vitamin B1, B2, and B6 Intake

Dietary intake of vitamins B1 (thiamine), B2 (riboflavin), and B6 (pyridoxine) was assessed using the two-day 24-hour dietary recall interviews collected in NHANES:

Day 1 Recall: Conducted via in-person interview using the Automated Multiple Pass Method (AMPM), a standardized protocol designed to reduce recall bias and improve data accuracy.

Day 2 Recall: Conducted by telephone to validate and enhance the reliability of dietary intake data.

The average intake of each vitamin over the two days was calculated for each participant. Dietary intake was further categorized into quartiles (Q1–Q4), with Q1 representing the lowest intake and Q4 the highest intake group.

### To ensure data quality

Extreme outliers (>3 standard deviations from the mean) were excluded. Average intake levels were visually examined for distribution patterns (Figure 2-4).

**Figure 2.**
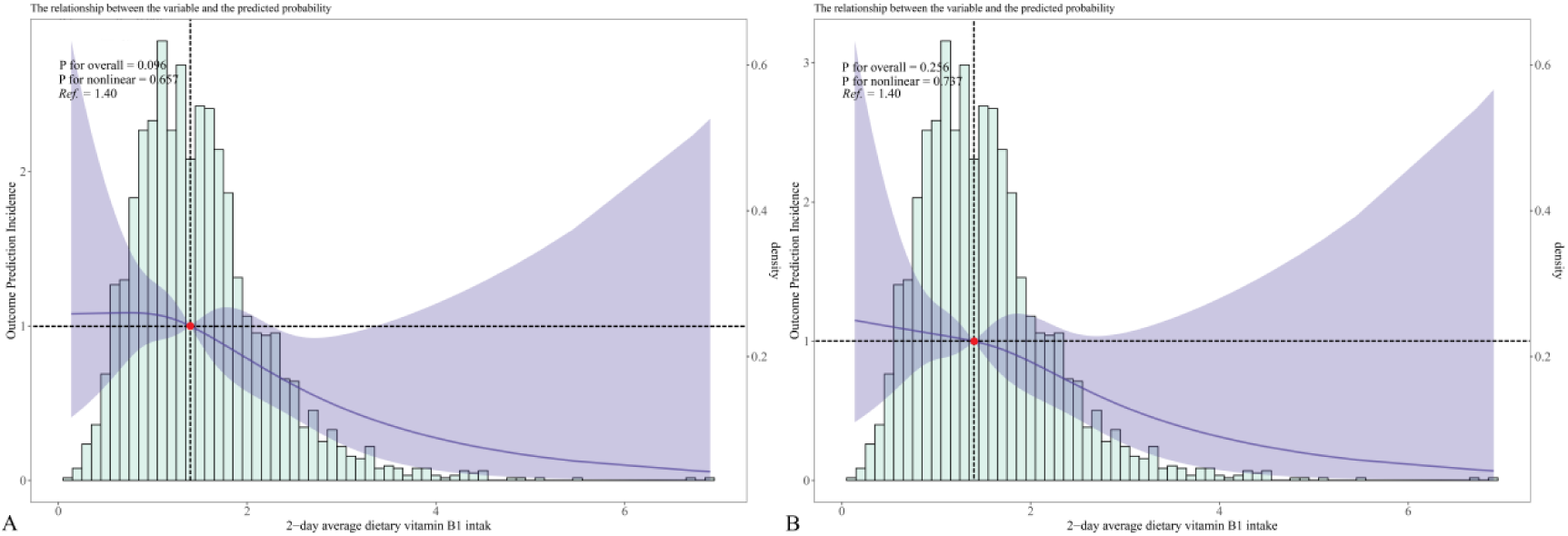
B1 RCS

**Figure 3.**
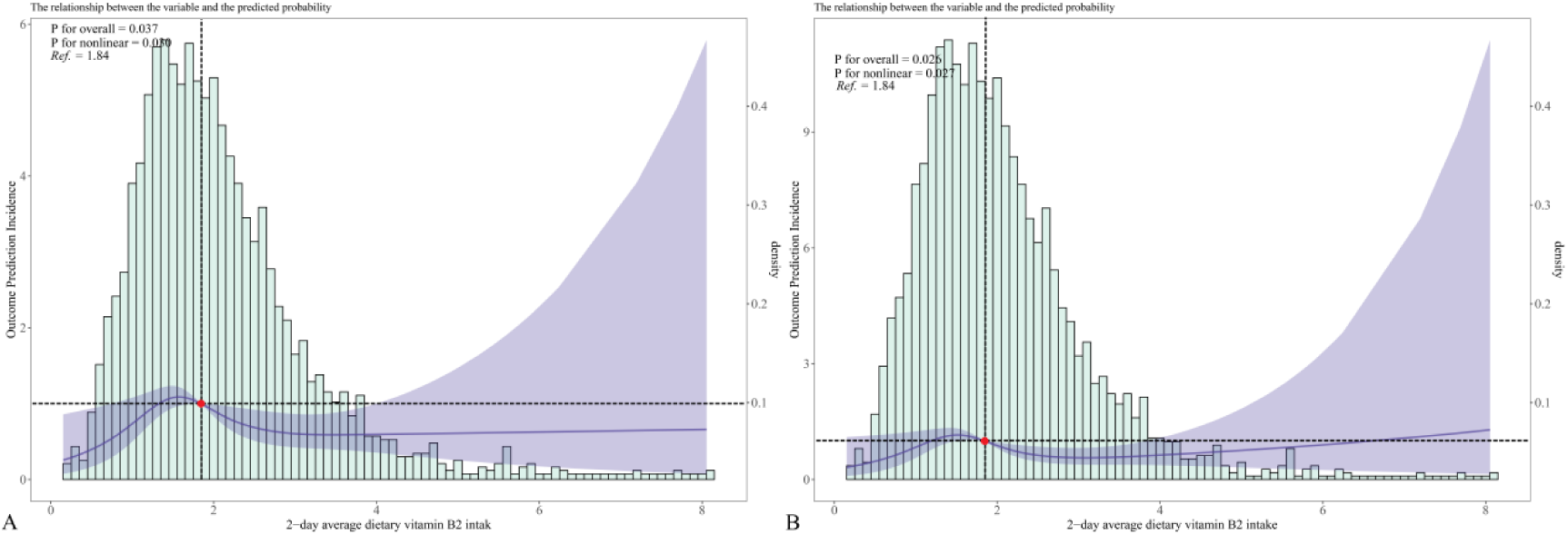
B2 RCS

**Figure 4.**
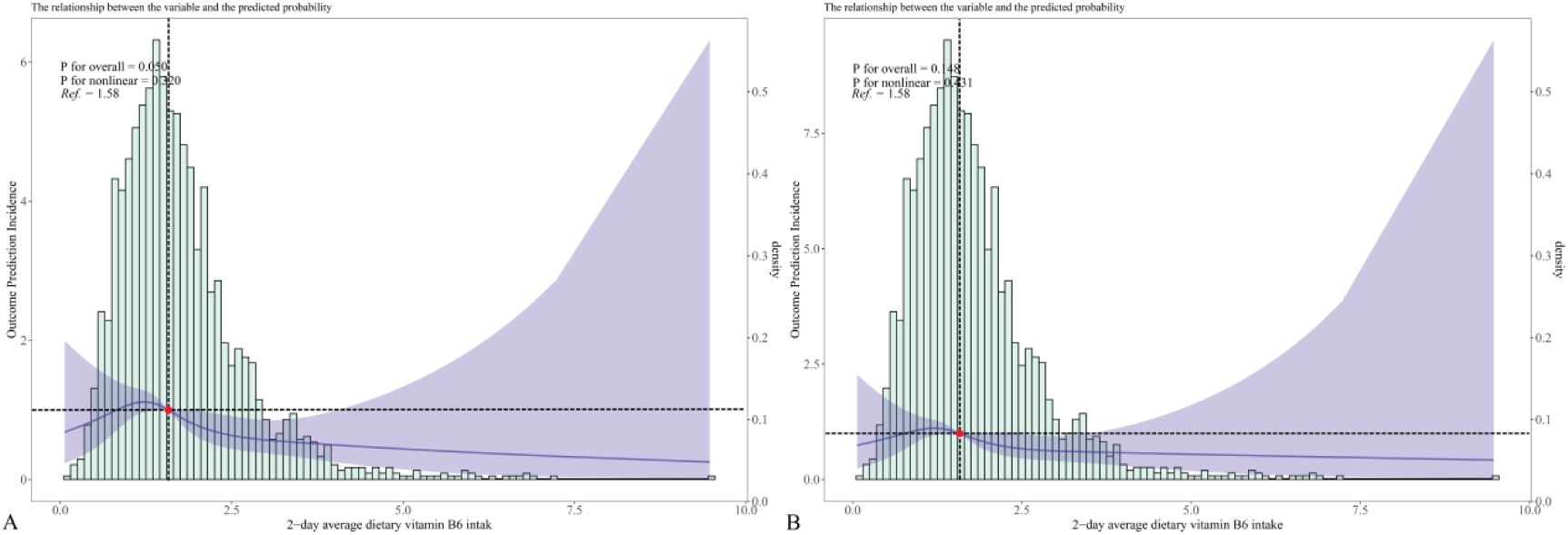
B6 RCS

### Outcome Assessment: Endometriosis

Endometriosis status was defined based on self-reported physician diagnosis, as obtained from the NHANES questionnaire:

“Has a doctor or other health professional ever told you that you have endometriosis?”

Participants who answered “Yes” were classified as cases, and those who answered “No” were classified as controls.

Although self-reported outcomes are subject to recall bias, prior studies have demonstrated acceptable validity for physician-diagnosed endometriosis in large epidemiological surveys.

### Covariates and Potential Confounders

Potential confounders were identified based on previous literature and biological plausibility. These covariates included:

Demographic Factors: Age (continuous, years), race/ethnicity (categorical: Non-Hispanic White, Non-Hispanic Black, Hispanic, Other Race), education level (<9th grade, high school diploma, college degree), and family income-to-poverty ratio (continuous).

Anthropometric Measures: BMI (continuous, kg/m²) and waist circumference (continuous, cm) were measured following standardized NHANES protocols.

Reproductive History: Menarche age (continuous, years), oral contraceptive use (yes/no), and pregnancy status (yes/no).

Lifestyle Factors: Smoking status (yes/no), drinking status (alcohol use: yes/no).

Clinical Factors: Hypertension status (yes/no).

The distribution of covariates between participants with and without endometriosis was assessed and summarized in Table 1.

**Table 1.** Characteristics of NHANES (2003–2006) participants with Endometriosis (N = 2462).

| Variables | Total (n = 2462) | No Endometriosis (n = 2296) | Endometriosis (n = 166) | *P* |
| --- | --- | --- | --- | --- |
| Age(year)<sup>a</sup>, Mean ± SD | 35.84 ± 10.09 | 35.51 ± 10.10 | 40.40 ± 8.75 | <.001 |
| Race/ethnicity<sup>b</sup>, n(%) |  |  |  | <.001 |
| Mexican American | 519 (21.08) | 505 (21.99) | 14 (8.43) |  |
| Other Hispanic | 93 (3.78) | 89 (3.88) | 4 (2.41) |  |
| Non-Hispanic White | 1190 (48.33) | 1082 (47.13) | 108 (65.06) |  |
| Non-Hispanic Black | 543 (22.06) | 508 (22.13) | 35 (21.08) |  |
| Other Race | 117 (4.75) | 112 (4.88) | 5 (3.01) |  |
| Education<sup>b</sup>, n(%) |  |  |  | 0.019 |
| <9th grade | 160 (6.50) | 156 (6.79) | 4 (2.41) |  |
| 9–11th grade | 348 (14.13) | 333 (14.50) | 15 (9.04) |  |
| High school diploma/GED | 532 (21.61) | 499 (21.73) | 33 (19.88) |  |
| Some College/AA degree | 844 (34.28) | 777 (33.84) | 67 (40.36) |  |
| ≥College graduate | 578 (23.48) | 531 (23.13) | 47 (28.31) |  |
| Family income to poverty ratio<sup>a</sup>, Mean ± SD | 2.68 ± 1.62 | 2.65 ± 1.62 | 3.21 ± 1.61 | <.001 |
| BMI<sup>a</sup>, Mean ± SD | 29.18 ± 7.60 | 29.21 ± 7.65 | 28.73 ± 6.73 | 0.438 |
| Waistline<sup>a</sup>, Mean ± SD | 95.69 ± 16.75 | 95.78 ± 16.87 | 94.44 ± 15.01 | 0.321 |
| Menarche age<sup>a</sup>, Mean ± SD | 12.54 ± 1.74 | 12.55 ± 1.73 | 12.33 ± 1.81 | 0.112 |
| Take contraceptive pills<sup>b</sup>, n(%) |  |  |  | <.001 |
| No | 562 (22.83) | 551 (24.00) | 11 (6.63) |  |
| Yes | 1900 (77.17) | 1745 (76.00) | 155 (93.37) |  |
| Pregnancy status<sup>b</sup>, n(%) |  |  |  | 0.001 |
| No | 2012 (81.72) | 1861 (81.05) | 151 (90.96) |  |
| Yes | 450 (18.28) | 435 (18.95) | 15 (9.04) |  |
| Hypertension<sup>b</sup>, n(%) |  |  |  | <.001 |
| No | 1996 (81.07) | 1879 (81.84) | 117 (70.48) |  |
| Yes | 466 (18.93) | 417 (18.16) | 49 (29.52) |  |
| Drinking status<sup>b</sup>, n(%) |  |  |  | 0.010 |
| No | 415 (16.86) | 399 (17.38) | 16 (9.64) |  |
| Yes | 2047 (83.14) | 1897 (82.62) | 150 (90.36) |  |
| Smoking status<sup>b</sup>, n(%) |  |  |  | 0.181 |
| No | 1772 (71.97) | 1660 (72.30) | 112 (67.47) |  |
| Yes | 690 (28.03) | 636 (27.70) | 54 (32.53) |  |
| Average dietary vitamin B1 intake<sup>a</sup>, Mean ± SD | 1.51 ± 0.71 | 1.52 ± 0.72 | 1.38 ± 0.59 | 0.003 |
| Average dietary vitamin B1 group<sup>b</sup>, n(%) |  |  |  | 0.149 |
| Q1 | 616 (25.02) | 568 (24.74) | 48 (28.92) |  |
| Q2 | 614 (24.94) | 567 (24.70) | 47 (28.31) |  |
| Q3 | 616 (25.02) | 575 (25.04) | 41 (24.70) |  |
| Q4 | 616 (25.02) | 586 (25.52) | 30 (18.07) |  |
| Average dietary vitamin B2 intake<sup>a</sup>, Mean ± SD | 2.01 ± 0.95 | 2.01 ± 0.96 | 1.93 ± 0.86 | 0.252 |
| Average dietary vitamin B2 group<sup>b</sup>, n(%) |  |  |  | 0.099 |
| Q1 | 616 (25.02) | 580 (25.26) | 36 (21.69) |  |
| Q2 | 615 (24.98) | 562 (24.48) | 53 (31.93) |  |
| Q3 | 615 (24.98) | 571 (24.87) | 44 (26.51) |  |
| Q4 | 616 (25.02) | 583 (25.39) | 33 (19.88) |  |
| Average dietary vitamin B6 intake<sup>a</sup>, Mean ± SD | 1.76 ± 0.92 | 1.77 ± 0.93 | 1.59 ± 0.79 | 0.008 |
| Average dietary vitamin B6 intake group<sup>b</sup>, n(%) |  |  |  | 0.032 |
| Q1 | 616 (25.02) | 566 (24.65) | 50 (30.12) |  |
| Q2 | 615 (24.98) | 575 (25.04) | 40 (24.10) |  |
| Q3 | 615 (24.98) | 566 (24.65) | 49 (29.52) |  |
| Q4 | 616 (25.02) | 589 (25.65) | 27 (16.27) |  |

a: Student t-test, b: Chi-square test, SD: standard deviation

### Statistical Analysis

All statistical analyses were performed using R version 4.1.3, accounting for NHANES’s complex sampling design (sampling weights, clusters, and strata) to produce nationally representative estimates.

#### Descriptive Analysis

Continuous variables (e.g., age, BMI, vitamin intake) were expressed as means ± standard deviations (SD) and compared using Student’s t-tests.

Categorical variables (e.g., race, oral contraceptive use) were expressed as percentages and compared using the chi-square test.

Correlation analysis among variables (e.g., BMI, waist circumference, vitamin intake, and socioeconomic factors) was visualized using a correlation matrix heatmap (Figure 5).

### Logistic Regression Models

Multivariate logistic regression models were applied to evaluate the association between dietary vitamin B1 intake (primary exposure) and the risk of endometriosis. Vitamins B2 and B6 served as secondary exposures for comparison. Odds ratios (ORs) and 95% confidence intervals (CIs) were reported for each model:

Model 1: Crude, unadjusted analysis.

Model 2: Adjusted for age, race/ethnicity, education, BMI, waist circumference, and family income-to-poverty ratio.

Model 3: Further adjusted for lifestyle factors (smoking and drinking status) and clinical factors (hypertension).

Model 4: Fully adjusted for reproductive factors (menarche age, pregnancy status, and oral contraceptive use).

To assess trends, the quartiles of vitamin intake were treated as an ordinal variable, and P-values for trends were calculated (Supplementary Table 1–3).

**Supplementary Table 1.** Trend analysis between 2-day average dietary vitamin B1 intake and the risk of endometriosis.

| Variables | Model1 |  | Model2 |  | Model3 |  | Model4 |  |
| --- | --- | --- | --- | --- | --- | --- | --- | --- |
|  | OR (95%CI) | P | OR (95%CI) | P | OR (95%CI) | P | OR (95%CI) | P |
| Trend Value | 0.71 (0.53 ~ 0.96) | 0.026 | 0.76 (0.55 ~ 1.03) | 0.080 | 0.76 (0.56 ~ 1.04) | 0.089 | 0.77 (0.56 ~ 1.06) | 0.105 |

OR: Odds Ratio, CI: Confidence Interval
Model1: Crude
Model2: Adjust: Age, Race/ethnicity, Education, Family income to poverty ratio, BMI, Waistline
Model3: Adjust: Age, Race/ethnicity, Education, Family income to poverty ratio, BMI, Waistline, Drinking status, Smoking status, Hypertension
Model4: Adjust: Age, Race/ethnicity, Education, Family income to poverty ratio, BMI, Waistline, Drinking status, Smoking status, Hypertension, Menarche age, Take contraceptive pills, Pregnancy status

**Supplementary Table 2.**
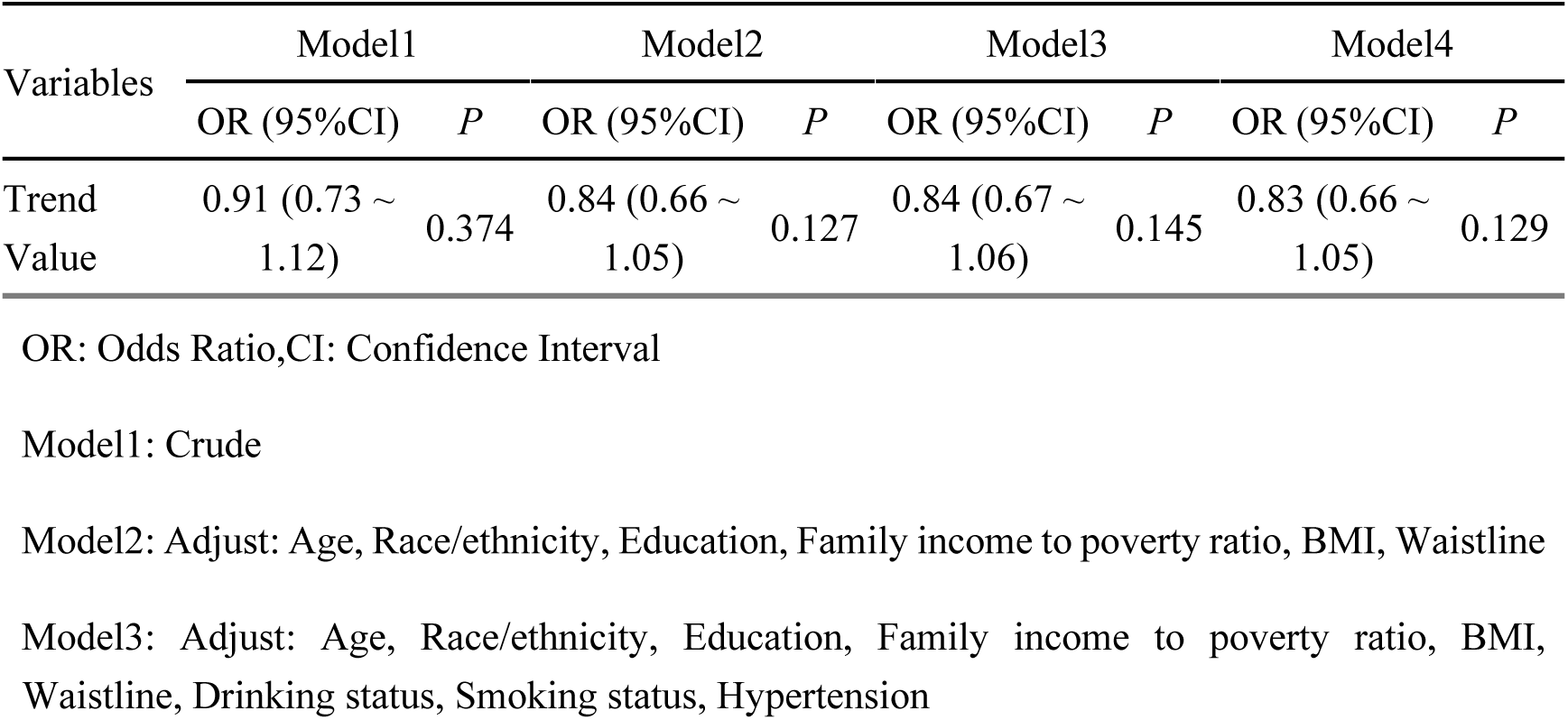
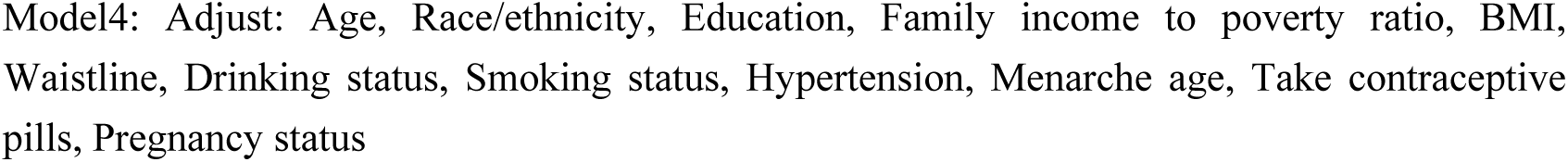
Trend analysis between 2-day average dietary vitamin B2 intake and the risk of endometriosis.

**Supplementary Table 3.**
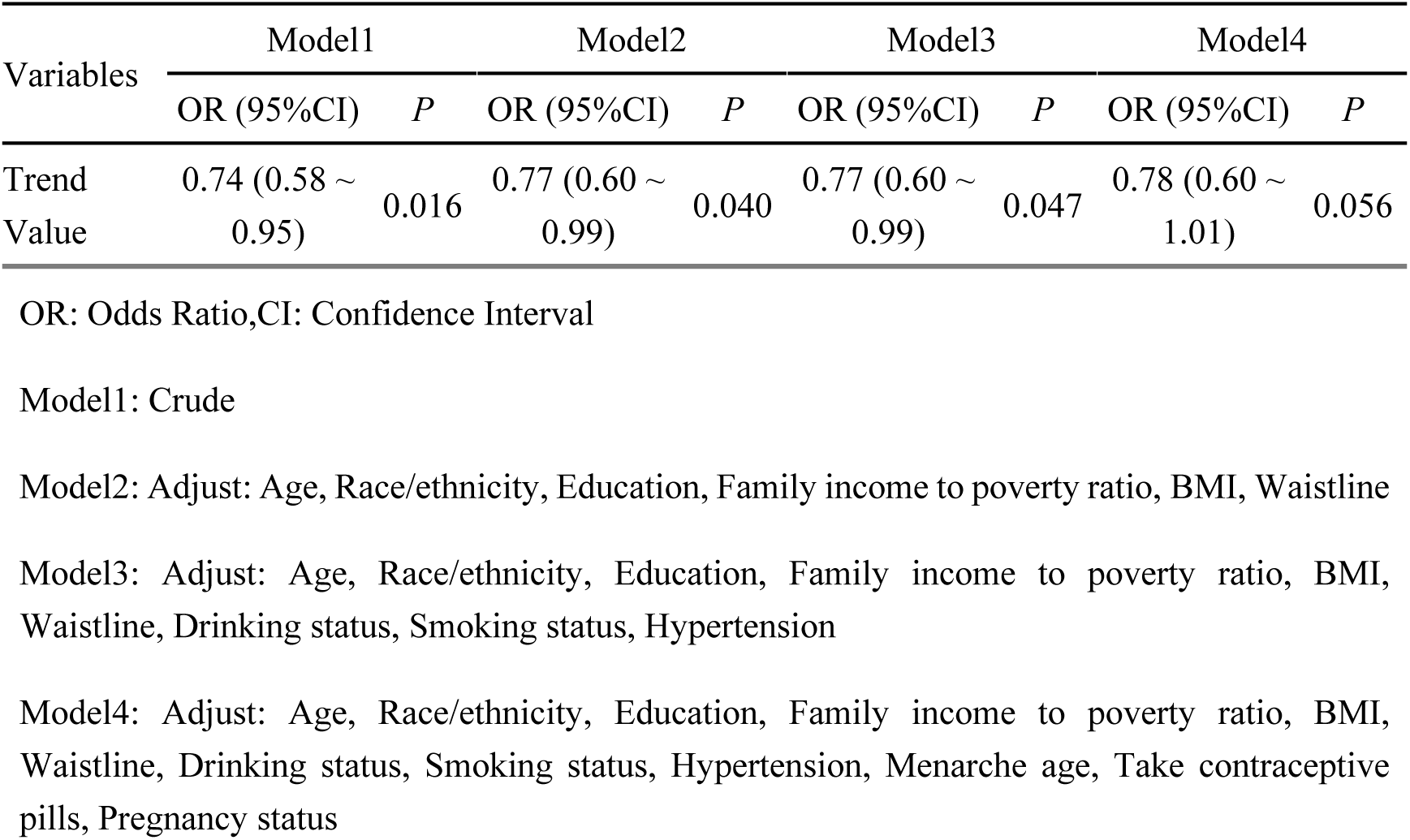
Trend analysis between 2-day average dietary vitamin B6 intake and the risk of endometriosis.

### Subgroup and Interaction Analysis

Stratified analyses were conducted to examine potential effect modifications by BMI (<25 vs. ≥25 kg/m²) and waist circumference (<93.7 cm vs. ≥93.7 cm) (Figures 6–8). Interaction terms between vitamin B1 intake and stratification variables were included in logistic models, and statistical significance was tested using likelihood ratio tests.

### Non-Linear Analysis

To investigate potential non-linear relationships between dietary vitamin B1 intake and endometriosis risk, restricted cubic spline (RCS) regression was performed with three knots (10th, 50th, and 90th percentiles). RCS analysis allowed identification of potential threshold effects and inflection points, where the relationship between vitamin B1 intake and endometriosis risk plateaued or changed direction (Figure 2).

### Sensitivity Analysis

Sensitivity analyses were conducted to ensure the robustness of the results:

Excluding participants with extreme vitamin B1 intake values.

Treating vitamin intake as a continuous variable instead of categorical quartiles.

### Data Visualization

A flowchart (Figure 1) illustrates the participant inclusion and exclusion process.

A correlation matrix (Figure 5) displays the inter-relationships among covariates.

**Figure 5.**
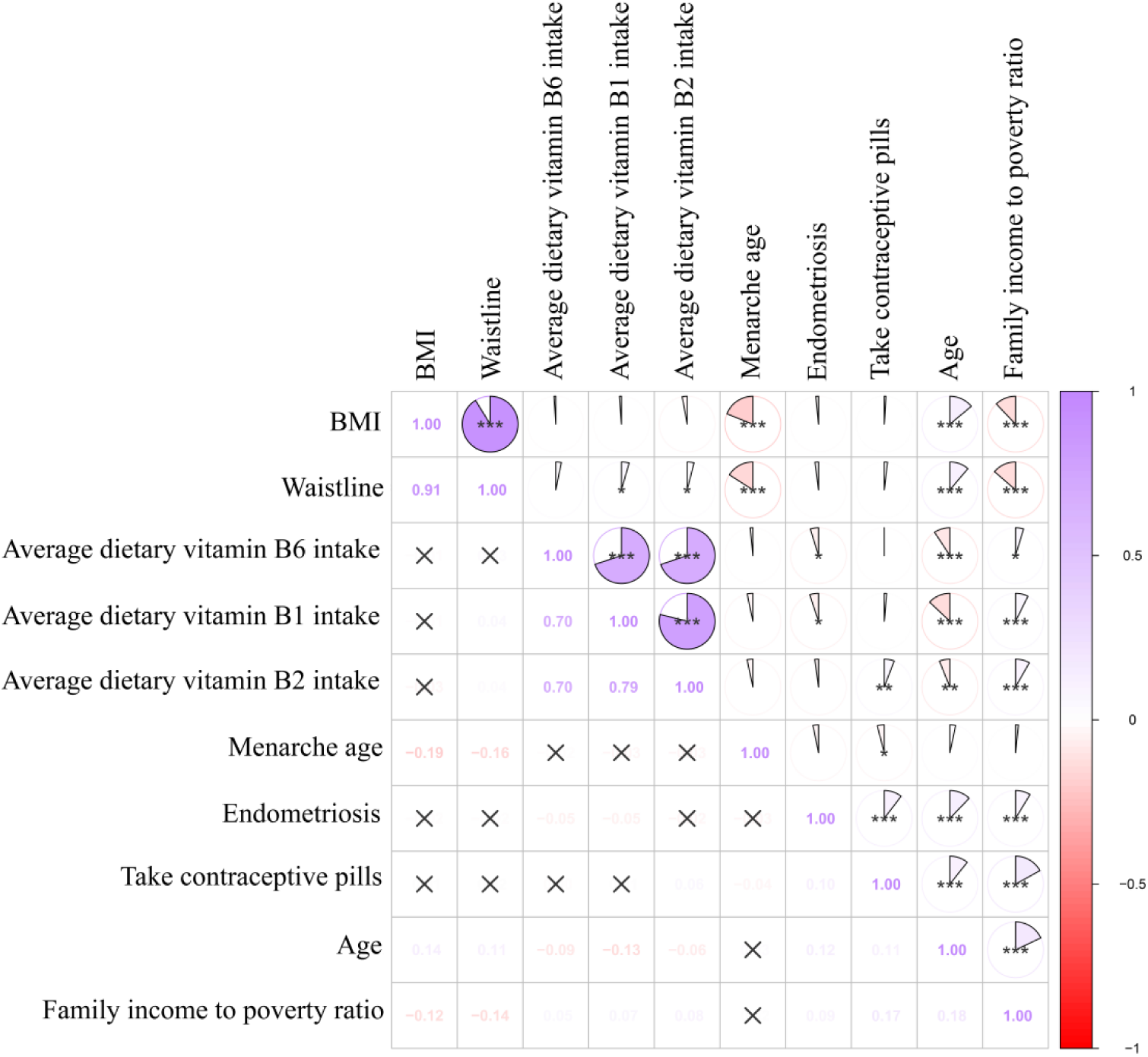
Correlation Matrix of Vitamin B1 Intake with BMI and Waist Circumference

Forest plots (Figures 6–8) depict subgroup analysis results, highlighting the protective effect of vitamin B1 intake in specific populations.

**Figure 6.**
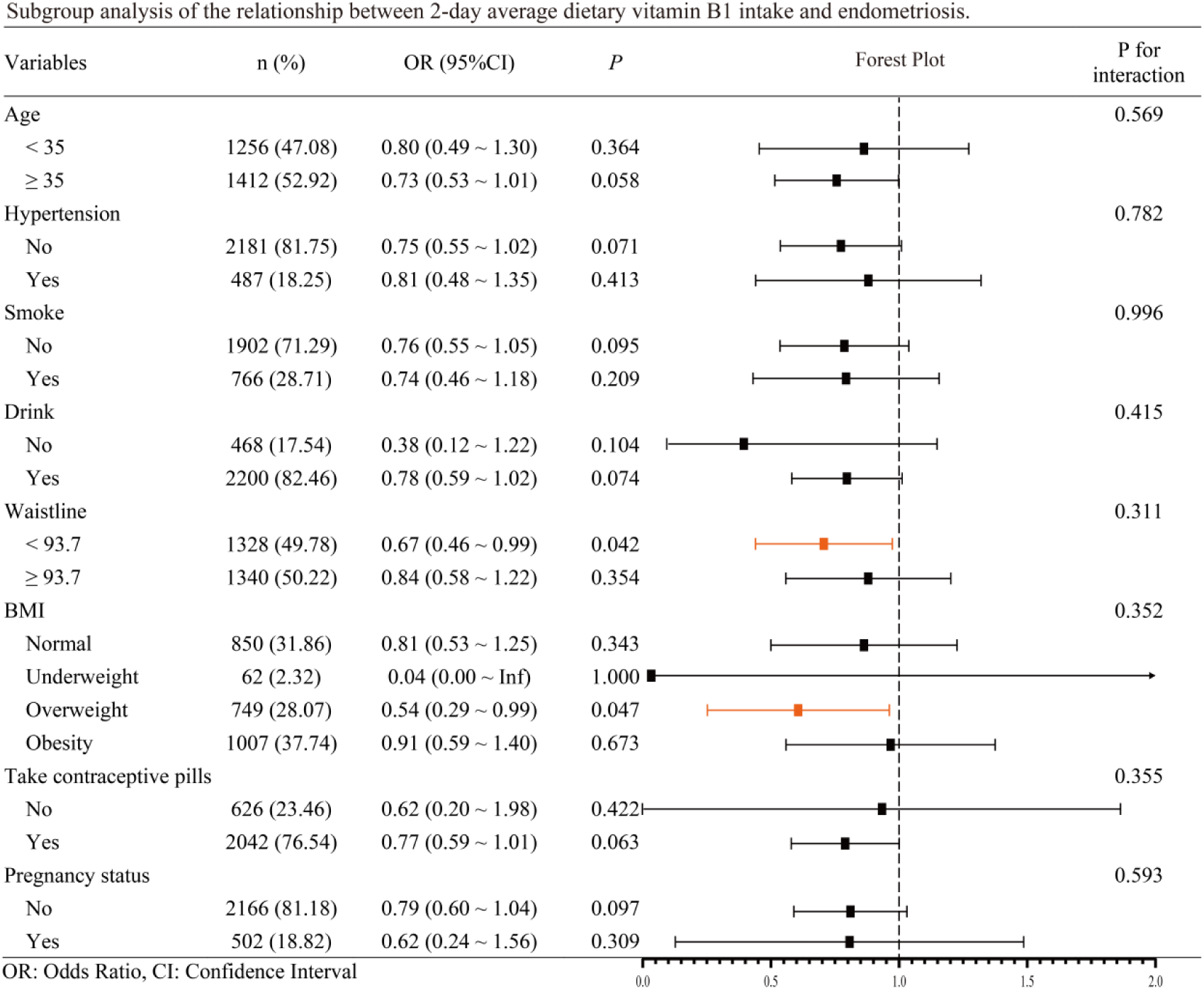
Subgroup Analysis for Vitamin B1

**Figure 7.**
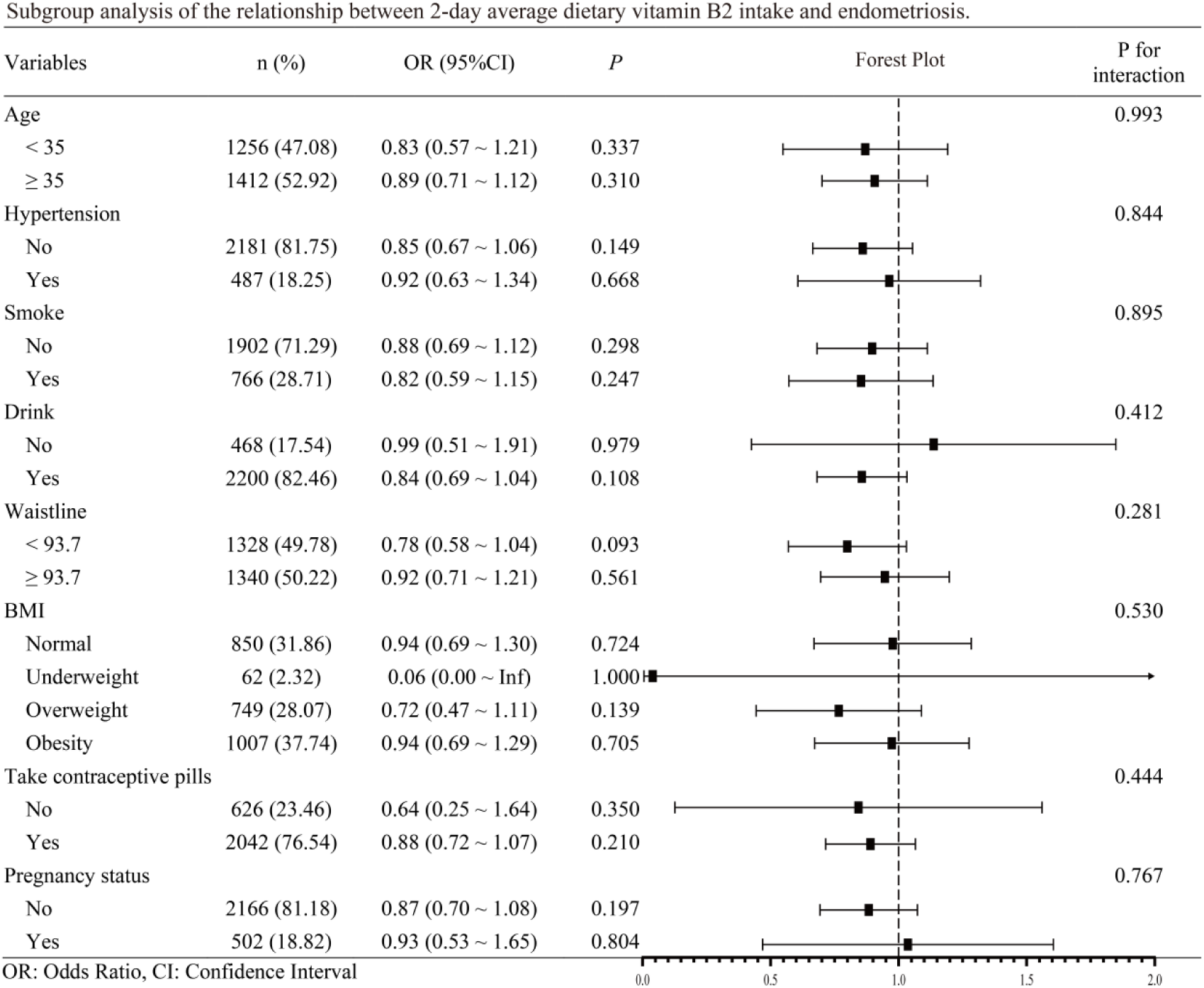
Subgroup Analysis for Vitamin B2

**Figure 8.**
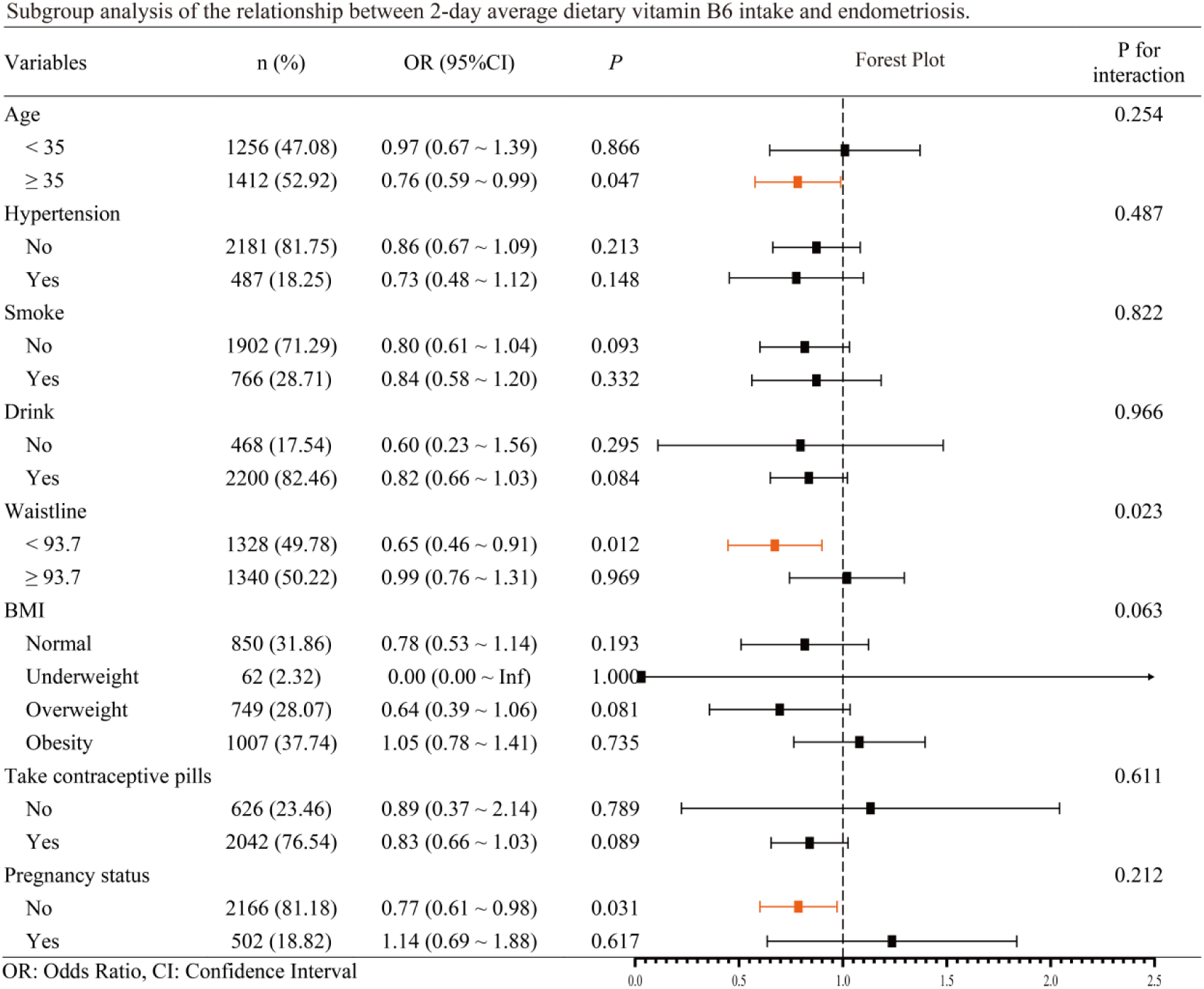
Subgroup Analysis for Vitamin B6

Trend analysis heatmaps (Figure 9) summarize odds ratios for all B vitamins across logistic models.

**Figure 9.**
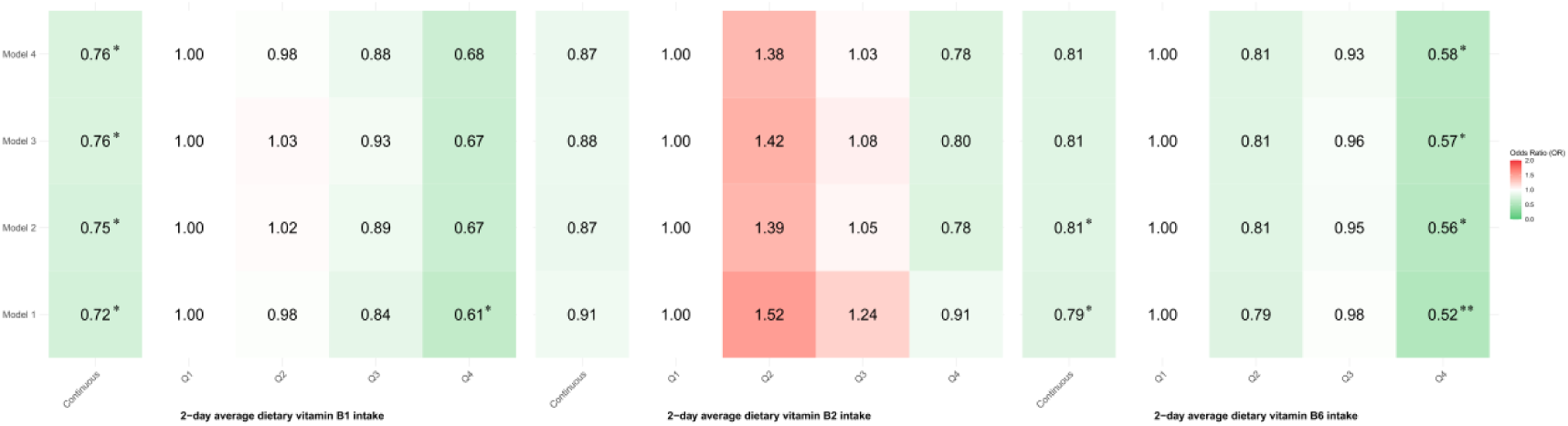
Heatmap Illustrating the Significant Protective Effect of Vitamin B1

Restricted cubic spline plots (Figures 2–4) visually represent non-linear relationships for vitamin B1, B2, and B6 intakes.

## Results

### Participant Characteristics

Baseline characteristics of the study population, stratified by endometriosis status, are summarized in Table 1. Among the 2,462 participants, 166 (6.7%) reported a diagnosis of endometriosis, while 2,296 (93.3%) did not. Key findings include:

Age: Participants with endometriosis were significantly older (40.4 ± 8.75 years) compared to those without endometriosis (35.5 ± 10.10 years, P < 0.001).

### Reproductive History

Menarche age was slightly younger in the endometriosis group (12.3 ± 1.81 years vs. 12.5 ± 1.73 years, P = 0.112).

Oral contraceptive use was markedly higher in the endometriosis group (93.4% vs. 76.0%, P < 0.001).

Anthropometric Measures: BMI and waist circumference were higher in participants with endometriosis, though not statistically significant (P = 0.438, P = 0.321, respectively).

### Dietary Intake

Vitamin B1 intake was significantly lower in participants with endometriosis (1.38 ± 0.59 mg/day) compared to controls (1.52 ± 0.72 mg/day, P = 0.003).

No significant differences were observed for vitamin B2 (P = 0.252).

Vitamin B6 intake showed a marginally significant difference (1.59 ± 0.79 mg/day vs. 1.77 ± 0.93 mg/day, P = 0.008).

These descriptive results indicate a potential protective effect of higher dietary vitamin B1 intake against endometriosis.

### Association Between Dietary Vitamin B1 Intake and Endometriosis Risk

Logistic regression analyses evaluated the relationship between vitamin B1 intake and endometriosis risk (Table 2):

**Table 2.**
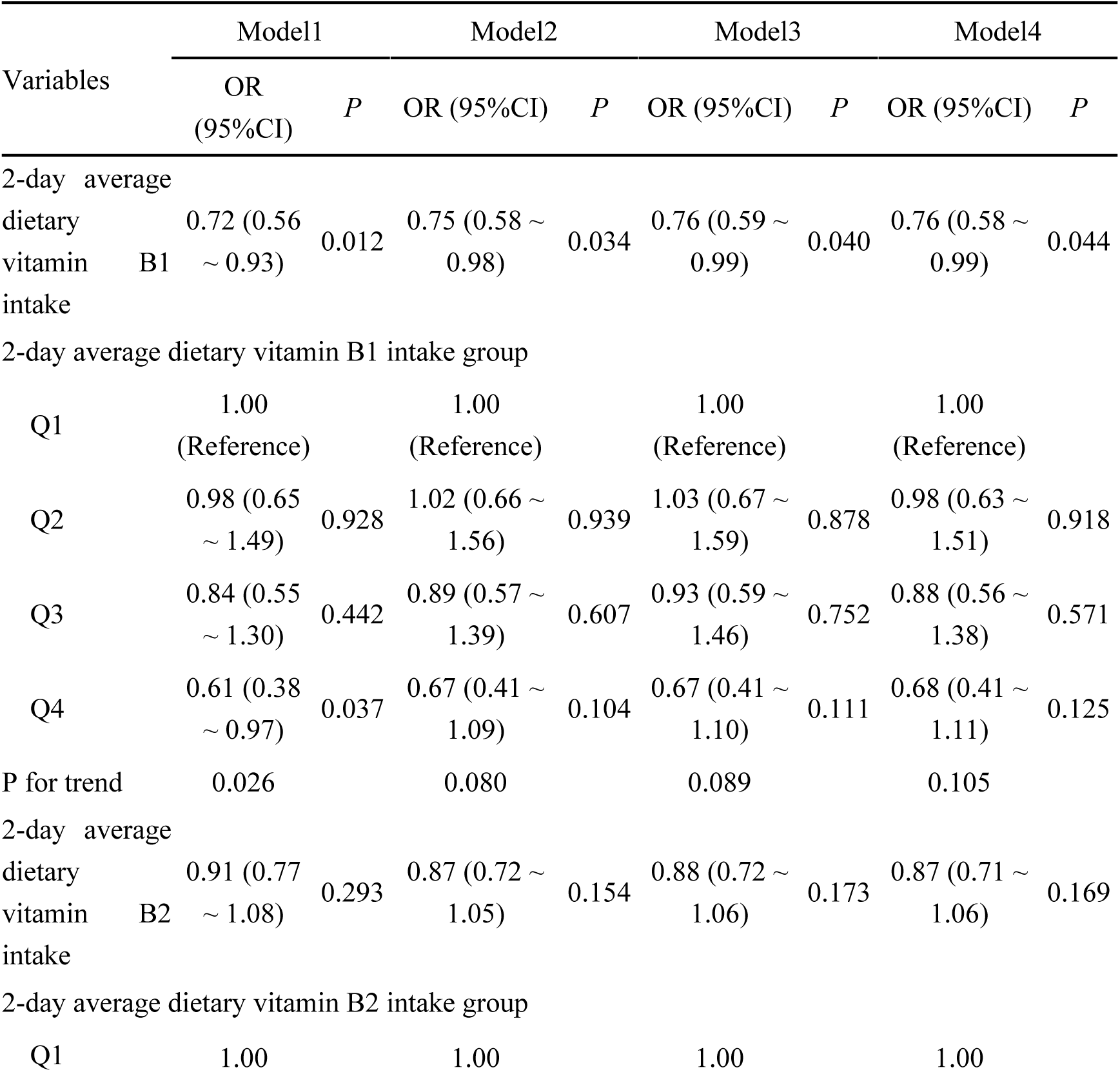
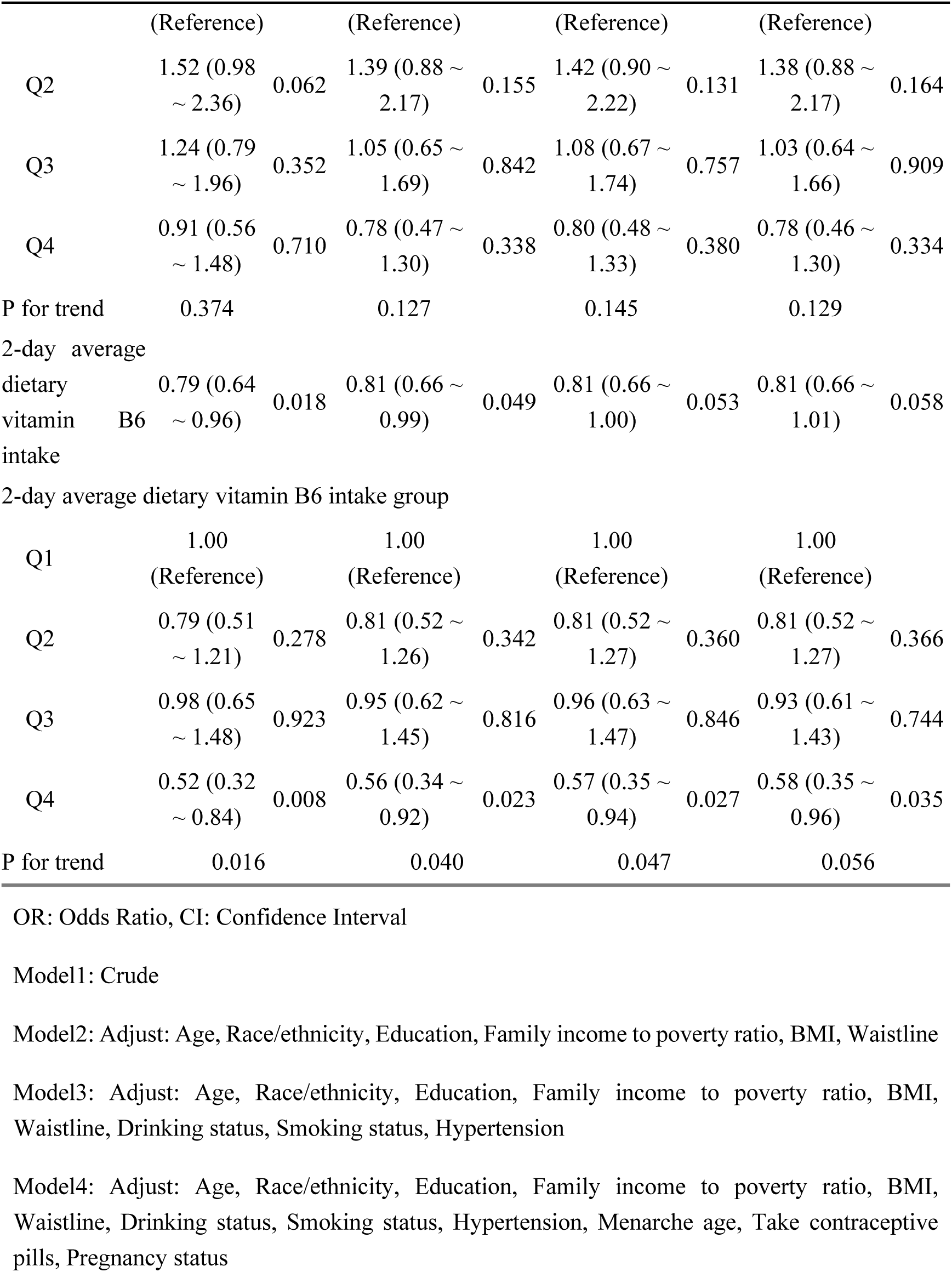
Association between different types of dietary vitamin B intake and the risk of endometriosis in NHANES (2003-2006).

Crude Model (Model 1): Higher dietary vitamin B1 intake was inversely associated with endometriosis risk (OR = 0.72, 95% CI: 0.56–0.93, P = 0.012).

Model 2: Adjusting for demographic covariates (age, race/ethnicity, education, BMI, waist circumference, and income) did not attenuate the association (OR = 0.75, 95% CI: 0.58–0.98, P = 0.034).

Model 3: Further adjustment for smoking status, drinking status, and hypertension yielded consistent results (OR = 0.76, 95% CI: 0.59–0.99, P = 0.040).

Model 4 (Fully Adjusted): After accounting for reproductive variables (menarche age, pregnancy status, and oral contraceptive use), the association remained significant (OR = 0.76, 95% CI: 0.58–0.99, P = 0.044).

Trend Analysis: As shown in Supplementary Table 1, a significant linear trend was observed across quartiles of vitamin B1 intake (P for trend = 0.026), with the highest quartile (Q4) demonstrating a 24% reduction in endometriosis risk compared to the lowest quartile (Q1).

### Comparative Analysis of Vitamins B2 and B6

In contrast to vitamin B1, the associations for vitamins B2 and B6 were weaker and not statistically significant (Supplementary Tables 2–3):

**Supplementary Table 2.**
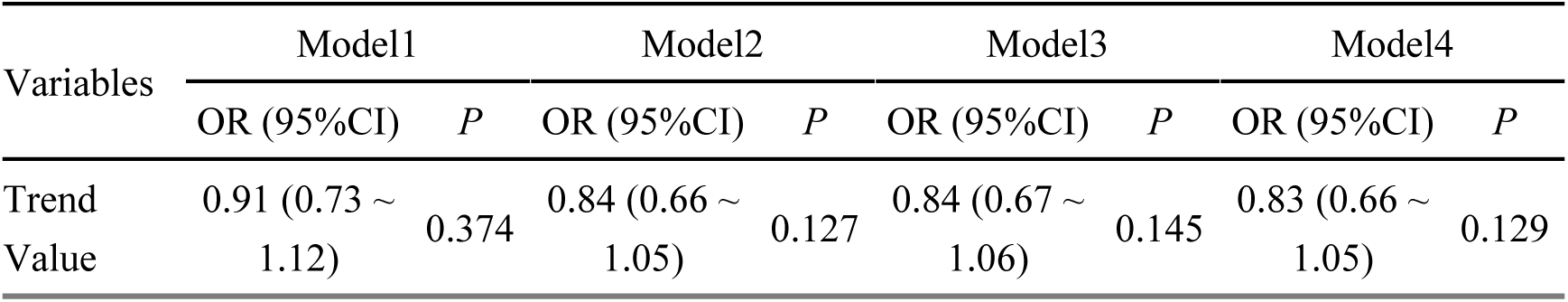
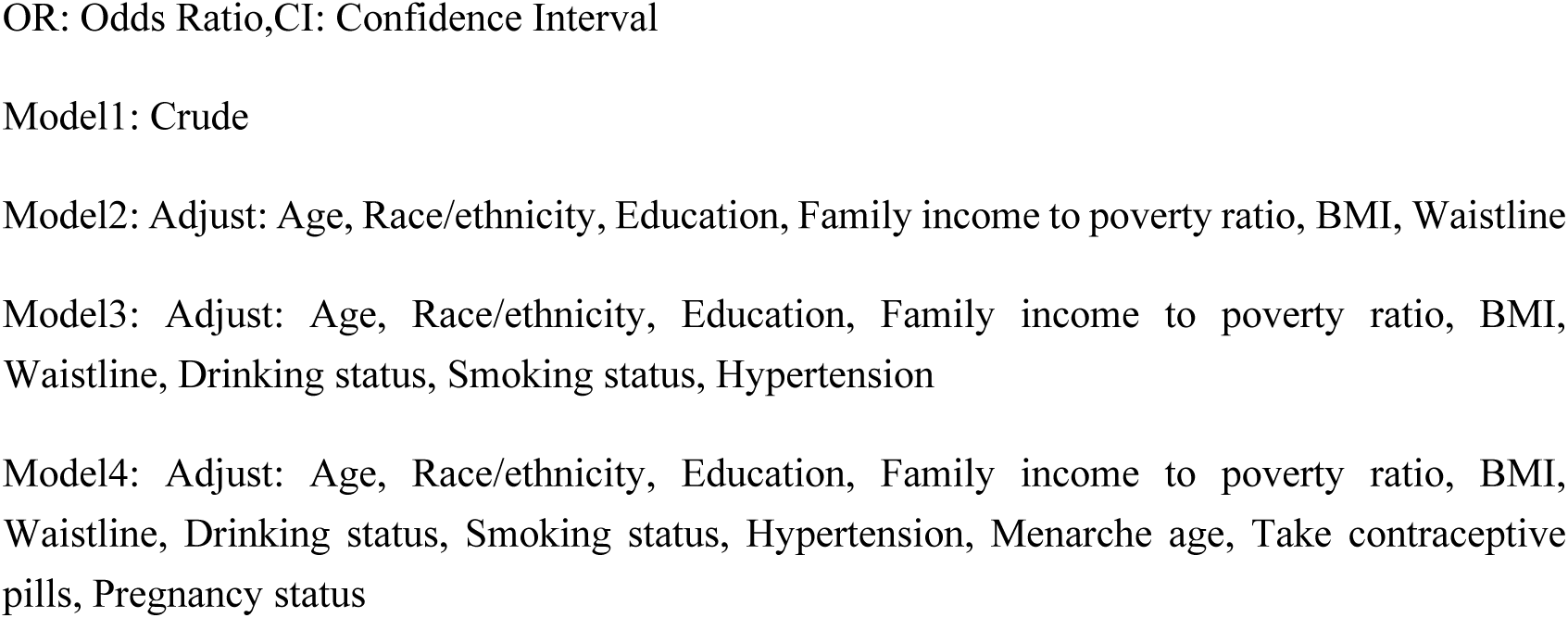
Trend analysis between 2-day average dietary vitamin B2 intake and the risk of endometriosis.

#### Vitamin B2

Model 4: OR = 0.83 (95% CI: 0.66–1.05, P = 0.129).

No significant linear trend across quartiles (P for trend = 0.127).

**Supplementary Table 3.** Trend analysis between 2-day average dietary vitamin B6 intake and the risk of endometriosis.

| Variables | Model1 |  | Model2 |  | Model3 |  | Model4 |  |
| --- | --- | --- | --- | --- | --- | --- | --- | --- |
|  | OR (95%CI) | P | OR (95%CI) | P | OR (95%CI) | P | OR (95%CI) | P |
| Trend Value | 0.74 (0.58 ~ 0.95) | 0.016 | 0.77 (0.60 ~ 0.99) | 0.040 | 0.77 (0.60 ~ 0.99) | 0.047 | 0.78 (0.60 ~ 1.01) | 0.056 |

OR: Odds Ratio, CI: Confidence Interval
Model1: Crude
Model2: Adjust: Age, Race/ethnicity, Education, Family income to poverty ratio, BMI, Waistline
Model3: Adjust: Age, Race/ethnicity, Education, Family income to poverty ratio, BMI, Waistline, Drinking status, Smoking status, Hypertension
Model4: Adjust: Age, Race/ethnicity, Education, Family income to poverty ratio, BMI, Waistline, Drinking status, Smoking status, Hypertension, Menarche age, Take contraceptive pills, Pregnancy status

#### Vitamin B6

Model 4: OR = 0.78 (95% CI: 0.60–1.01, P = 0.056).

A marginally significant trend was noted (P for trend = 0.056).

These results suggest that while vitamin B2 and B6 may have potential protective effects, their associations are not as robust as that of vitamin B1.

### Subgroup Analysis

To assess effect modifications, subgroup analyses were performed by BMI and waist circumference (Figures 6–8):

#### BMI Stratification

Participants with BMI < 25 demonstrated a stronger inverse association (OR = 0.54, 95% CI: 0.29–0.99, P = 0.047).

No significant association was observed in participants with BMI ≥ 25 (P > 0.05).

#### Waist Circumference Stratification

Participants with waist circumference < 93.7 cm exhibited a significant reduction in endometriosis risk (OR = 0.67, 95% CI: 0.46–0.99, P = 0.042).

No significant association was observed in participants with waist circumference ≥ 93.7 cm.

These findings suggest that the protective effect of vitamin B1 intake is more prominent among women with normal weight and waist circumference.

### Non-Linear Analysis of Vitamin B1 Intake

Restricted cubic spline regression (RCS) was used to explore the non-linear relationship between vitamin B1 intake and endometriosis risk (Figure 2):

A significant non-linear relationship was observed (P for non-linearity = 0.037).

The risk of endometriosis decreased steadily with increasing vitamin B1 intake up to an inflection point of 1.84 mg/day. Beyond this point, the protective effect plateaued.

For comparison, RCS analyses for vitamins B2 and B6 (Figures 3–4) showed no significant non-linear trends (P > 0.05).However, the detailed relationship between dietary vitamin B1 intake and predicted probability is visually depicted in Figure 9.

### Correlation Analysis

The relationships between dietary intake and covariates were visualized using a correlation matrix (Figure 5):

BMI and waist circumference were strongly positively correlated (r = 0.91).

Vitamin B1 intake was weakly negatively correlated with BMI (r = −0.19) and waist circumference (r = −0.16), consistent with the subgroup analysis results.

No strong correlations were observed for vitamin B2 or B6 intake with other covariates.

### Sensitivity Analyses

To verify the robustness of the findings, sensitivity analyses were performed:

Excluding extreme outliers in vitamin intake values did not alter the results (data not shown).

Treating vitamin B1 intake as a continuous variable yielded consistent inverse associations with endometriosis risk (P < 0.05).

## Discussion

This study investigated the association between dietary vitamin B1 intake and the risk of endometriosis in a nationally representative cohort from NHANES 2003–2006. The results revealed a significant inverse association between dietary vitamin B1 intake and the risk of endometriosis, particularly among women with normal BMI and waist circumference. Vitamins B2 and B6, included as reference nutrients, showed weaker or non-significant associations, suggesting a specific protective role of vitamin B1. The discussion elaborates on these findings in light of existing evidence, plausible biological mechanisms, and potential clinical and public health implications.

### Primary Findings and Comparison With Previous Studies

The inverse association between dietary vitamin B1 intake and endometriosis risk aligns with prior research on the role of B vitamins in reducing oxidative stress and inflammation, two critical contributors to the pathogenesis of endometriosis ^5^; ^14^; ^15^. Vitamin B1, as a coenzyme in the pentose phosphate pathway, plays a key role in generating NADPH, which is essential for maintaining cellular redox balance ^16^. A higher dietary intake of vitamin B1 may enhance antioxidant capacity, thereby mitigating the inflammatory and oxidative damage observed in the peritoneal environment of women with endometriosis ^12^; ^17^.

Previous studies have predominantly focused on broader dietary patterns or specific anti-inflammatory nutrients. For instance, a study by Parazzini et al. ^5^ highlighted the role of antioxidant-rich foods, such as fruits and vegetables, in reducing endometriosis risk. However, few studies have specifically examined vitamin B1. In a related context, Harris et al. ^18^ reported that B vitamins, particularly B6, reduced systemic inflammation in reproductive-aged women, although their study did not directly address endometriosis. The findings from the present study extend this knowledge by identifying vitamin B1 as a potentially protective factor, independent of other B vitamins.

### Biological Mechanisms Underpinning Vitamin B1’s Protective Effect

The protective role of vitamin B1 can be attributed to its critical involvement in cellular energy metabolism and its antioxidant properties. Vitamin B1 functions as a coenzyme for transketolase, a key enzyme in the pentose phosphate pathway, which generates NADPH. NADPH is essential for the regeneration of reduced glutathione, a major intracellular antioxidant ^19^.

In endometriosis, oxidative stress is thought to drive chronic inflammation and promote ectopic implantation and proliferation of endometrial tissue ^20^; ^21^. Studies have shown elevated levels of reactive oxygen species (ROS) in the peritoneal fluid of women with endometriosis, along with re,duced antioxidant capacity ^22^; ^23^. By enhancing the cellular antioxidant defense system, vitamin B1 may counteract these pathological processes, thereby reducing the risk or progression of endometriosis.

The non-linear relationship observed in this study, with an inflection point at 1.84 mg/day of dietary B1 intake, suggests that optimal intake levels may confer maximal protection. This finding is consistent with previous research on the threshold effects of micronutrients in disease prevention ^24^.

### Subgroup Findings: BMI and Waist Circumference

Subgroup analyses revealed that the protective effect of vitamin B1 intake was more pronounced in women with BMI < 25 and waist circumference < 93.7 cm. These findings may be explained by differences in systemic inflammation and metabolic health across BMI categories. Obesity and central adiposity are associated with higher levels of pro-inflammatory cytokines and oxidative stress markers ^25^. This heightened inflammatory state may overwhelm the protective effects of dietary antioxidants, such as vitamin B1, in women with higher BMI or waist circumference ^26^.

These results underscore the importance of considering body composition in dietary interventions for endometriosis prevention. They also align with studies demonstrating stronger protective effects of antioxidant-rich diets in individuals with lower baseline levels of inflammation ^27^.

### Comparison With Vitamins B2 and B6

While vitamin B2 and B6 are also involved in antioxidant and anti-inflammatory pathways, their associations with endometriosis risk were weaker or non-significant in this study. Vitamin B2, as a cofactor for glutathione reductase, contributes to the recycling of oxidized glutathione, but its effects may be limited compared to the more direct NADPH-regenerating role of vitamin B1 ^28^.

Vitamin B6 has well-documented anti-inflammatory properties through its role in modulating cytokine production and homocysteine metabolism ^29^. However, its borderline significance in this study may reflect lower intake levels or insufficient statistical power to detect significant effects. Further research is needed to clarify the roles of these vitamins in endometriosis pathogenesis.

### Public Health Implications

The findings of this study have several public health and clinical implications. First, dietary vitamin B1 intake may represent a modifiable risk factor for endometriosis, particularly among women with normal BMI and waist circumference. Promoting vitamin B1-rich foods, such as whole grains, legumes, and lean meats, could be a cost-effective strategy for reducing the burden of this condition ^18^; ^22^; ^30^.

Second, the non-linear relationship between B1 intake and endometriosis risk highlights the importance of achieving optimal intake levels. This has implications for dietary guidelines and supplementation strategies, particularly for women at higher risk of endometriosis.

Finally, the findings underscore the need for prospective studies to confirm these associations and explore the potential for dietary interventions to complement existing medical and surgical treatments for endometriosis.

### Strengths and Limitations

#### Strengths

Use of a nationally representative sample from NHANES enhances the generalizability of the findings.

Comprehensive adjustment for demographic, lifestyle, and reproductive confounders ensures robust results.

Advanced statistical methods, including restricted cubic spline regression, provide insights into non-linear relationships.

#### Limitations

The cross-sectional design precludes causal inference, necessitating confirmation in longitudinal studies.

Dietary intake was self-reported and subject to recall bias, though the two-day dietary recall method employed by NHANES is well-validated. Self-reported endometriosis diagnosis may introduce misclassification bias, as some cases may be undiagnosed or inaccurately reported.

### Future Directions

#### Future research should focus on

Prospective cohort studies to establish causal relationships between dietary vitamin B1 intake and endometriosis risk.

Mechanistic studies to elucidate the specific pathways through which vitamin B1 modulates oxidative stress and inflammation in endometriosis.

Clinical trials to evaluate the efficacy of dietary interventions or supplementation strategies targeting B vitamins in preventing or managing endometriosis.

## Conclusion

This study provides novel evidence for the inverse association between dietary vitamin B1 intake and the risk of endometriosis, with the protective effect being particularly evident among women with normal BMI and waist circumference. In contrast, vitamins B2 and B6 showed weaker or non-significant associations, highlighting the unique role of vitamin B1 in modulating oxidative stress and inflammation, two key mechanisms underlying the pathogenesis of endometriosis.

The findings underscore the potential of dietary vitamin B1 as a modifiable risk factor for endometriosis prevention. Public health interventions promoting vitamin B1-rich foods, such as whole grains, legumes, and lean meats, could serve as a cost-effective strategy to reduce the burden of endometriosis. Furthermore, the observed non-linear relationship suggests that achieving optimal vitamin B1 intake levels is critical for maximizing its protective effects.

However, this cross-sectional study cannot establish causality, and further prospective cohort studies and clinical trials are warranted to confirm these findings and explore the efficacy of dietary interventions or supplementation strategies. Future research should also investigate the underlying biological mechanisms through which vitamin B1 influences endometriosis risk and examine its potential synergistic effects with other micronutrients.

In conclusion, this study highlights dietary vitamin B1 as a promising target for endometriosis prevention, contributing to a growing body of evidence on the role of nutrition in women’s reproductive health.

The data presented in this manuscript or supplementary information file supports the findings of this study and can be obtained from NHANES (National Health and Nutrition Examination Survey).The detailed data supporting the findings of this study are available from the corresponding author upon request.

## Data availability

The data presented in this paper can be obtained from National Health and Nutrition Examination Survey (NHANES, which can be accessed from https://www.cdc.gov/nchs/nhanes/index.htm). If you have any questions about the data or supplementary information files that support the findings of this study, please contact.

## Acknowledgements

We would like to express our gratitude to China, Jiangxi Province, and its affiliated institutions for providing funding for this research. Dr. Zhou Zhigang and Dr. Guo Huiwen are gratefully for their support of this study.

## Author Contributions Statement

Author Contributions Statement J. Z. and Z. Z. designed the study and performed data analysis. J. Z.collected the experimental data and conducted preliminary analyses. J. Z. was responsible for writing the main manuscript text. J. Z. and H. G. prepared Figures 1-9. All authors reviewed and edited the manuscript and approved the final version for submission. J. Z. and Z. Z. had full access to all the data in the study and takes responsibility for the integrity of the data and the accuracy of the data analysis. Z. Z. is the guarantor.

## Additional information

Publisher’s note

Springer Nature remains neutral with regard to jurisdictional claims in published maps and institutional affiliations.

## Clinical trial number: not applicable

This study utilized data from the NHANES, which is a publicly available resource. Data were fully anonymized, and no identifiable information was accessed. Ethical approval was not required according to the database’s terms of use and local regulations.

## Human Ethics and Consent to Participate declarations

not applicable

